## Supplemental information for "The transcriptional landscape of Venezuelan equine encephalitis virus (TC-83) infection"

### Supporting Information

**S1 Text. Rare structural viral read variants correlate with expression of specific host genes.** The high coverage corresponding to almost the entire viral genome obtained via viscRNA-Seq enabled further investigation of the structure of viral reads. Among millions of viral reads detected, we observed 14,956 gap reads (~0.1% of total viral reads), defined by having a deletion within read 1 or read 2 (we used Illumina paired-end sequencing, see Methods), not including reads with gaps between the two reads (**S3A Fig**). These gap reads were present in 271 cells and their abundance strongly correlated with vRNA abundance in the same cells, indicating that deep viral coverage was required for detection (**S3B Fig**). The length of these gaps ranged from 20 to over 10,000 nucleotides, with the majority being shorter than 1000 nucleotides (**S3C Fig**). The most common was a 36-base gap located within the coding region of the 6K protein (black arrow in Fig S3A). This gap was found in a total of 1,226 reads derived from 55 different cells. Prediction of the RNA structure via RNAfold web server (1) revealed that in the presence of the 36-base gap, there is formation of a hairpin with a free energy of -21.23 kcal/mol, indicating a very stable structure (**S3D Fig**). Although the biological function of this hairpin is unknown, stable RNA structures play essential roles in viral replication and tropism across multiple viruses. Alternatively, we cannot currently exclude that this gap could be a result of polymerase errors during the library preparation.

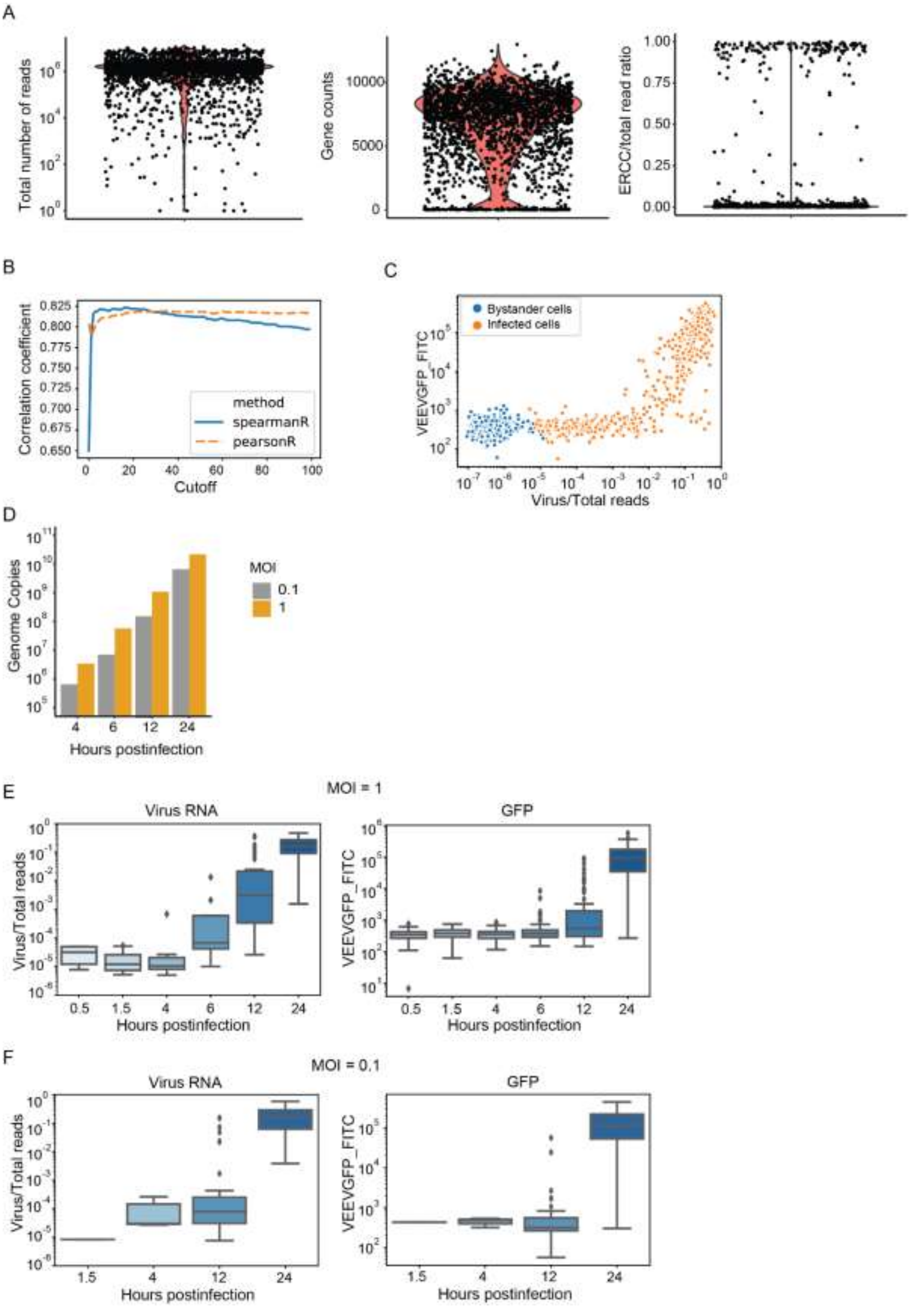
A
Total number of reads
Gene counts
ERCC/total read ratio
B
Correlation coefficient
method
spearmanR
pearsonR
Cutoff
C
Bystander cells
Infected cells
VEEVGFP_FITC
Virus/Total reads
D
Genome Copies
MOI
0.1
1
Hours postinfection
E
MOI = 1
Virus RNA
GFP
Virus/Total reads
VEEVGFP_FITC
Hours postinfection
F
MOI = 0.1
Virus RNA
GFP
Virus/Total reads
VEEVGFP_FITC
Hours postinfection

**S1 Fig. Quality control and definition of infected cells**. (A) Quality control of the VEEV-TC-83 infected cells dataset. Shown are the total number of reads (left panel), gene counts (middle panel) and ratio of ERCC spike-in RNA reads to total reads (right panel). Cell selection criteria included: total reads > 300,000, gene counts > 500 and a ratio of ERCC spike-in RNA to total reads < 0.05. (B) Spearman's and Pearson correlation coefficients between GFP expression and vRNA reads when using different cutoffs (from 0 to 100) to define infected cells. (C) Scatter plots showing GFP expression level and vRNA/total reads in cells with detectable vRNA reads. Cells harboring more than 10 vRNA reads. Orange, infected cells; blue, bystander cells. (D) Genome copies of VEEV-TC-83 per 500 ng total RNA at various time points postinfection at MOI 0.1 or 1. (E and F) Box plots showing virus/total read ratio (left panel) and GFP expression level (right panel) over time in infected cells (viral reads > 10) at an MOI of 1 (E) or 0.1 (F) (no infected cells were detected at an MOI of 0.1 at 0.5 and 6 hpi). HPI, hours postinfection; MOI, multiplicity of infection; ERCC, External RNA Controls Consortium.

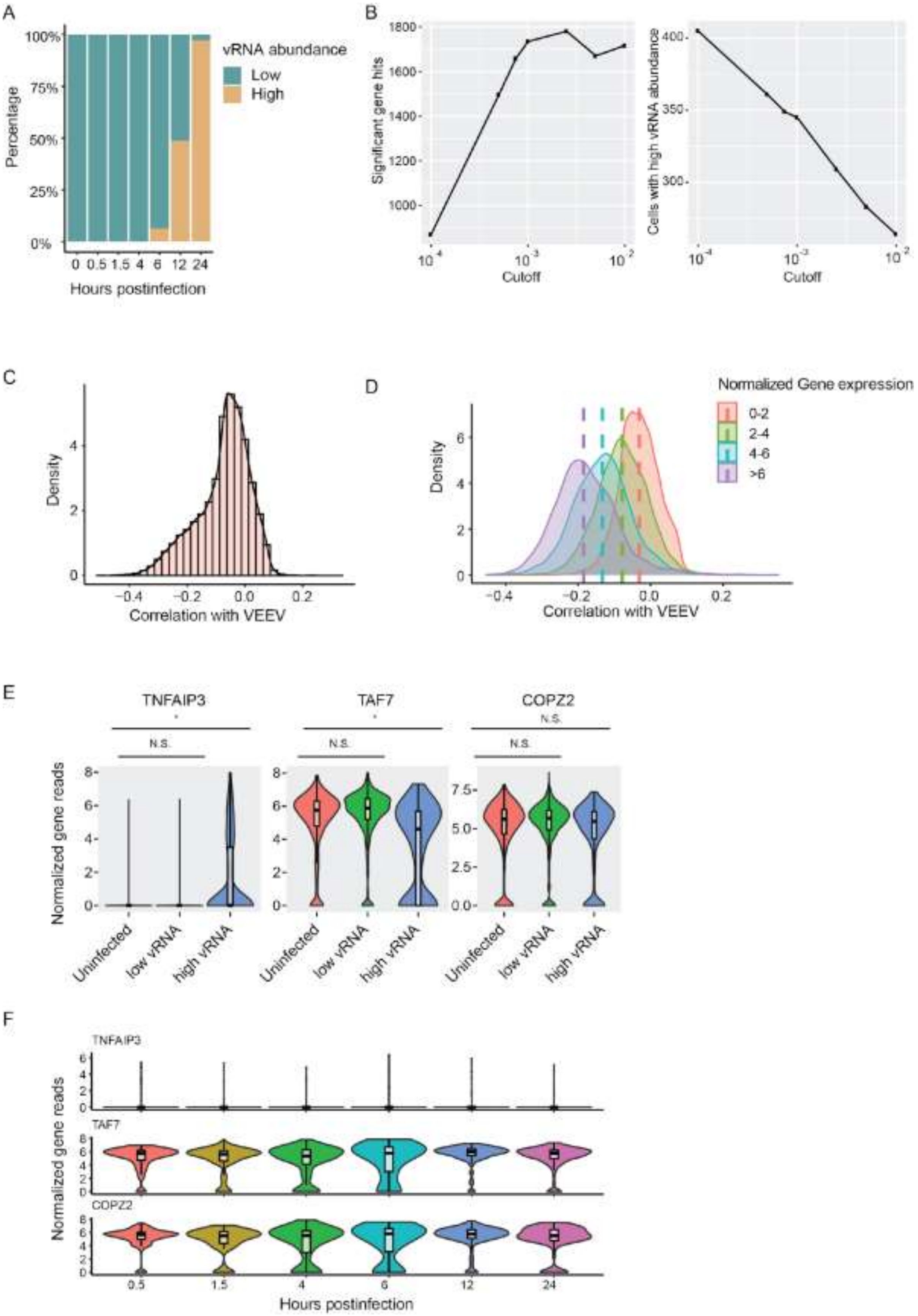
A
vRNA abundance
Low
High
Percentage
100%
75%
50%
25%
0%
0 0.5 1.5 4 6 12 24
Hours postinfection
B
Significant gene hits
Cells with high vRNA abundance
Cutoff
C
Density
Correlation with VEEV
D
Normalized Gene expression
0-2
2-4
4-6
>6
Density
Correlation with VEEV
E
TNFAIP3
TAF7
COPZ2
N.S.
Normalized gene reads
Uninfected
low vRNA
high vRNA
F
TNFAIP3
TAF7
COPZ2
Normalized gene reads
0.5 1.5 4 6 12 24
Hours postinfection

**S2 Fig. Subgrouping cells based on viral load, representative differentially expressed genes (DGEs) and correlation analysis**. (A) Percentage of low and high vRNA-harboring cells at each time point. High vRNA-harboring cells are defined as virus cDNA reads/total cDNA reads > 0.001. (B) The number of differentially expressed genes (left panel) and cells with high vRNA abundance (right panel) under different cutoffs set by a range of virus cDNA/total reads (from 0.0001 to 0.01). (C) Distribution of Spearman's correlation coefficients between VEEV-TC-83 vRNA abundance and ~55,000 host genes. (D) Distributions of Spearman's correlation coefficients shown in D stratified by the average expression level of the gene in uninfected cells. N.S., not significant; (E) Representative genes with distinct expression patterns between uninfected, low vRNA and high vRNA cell groups. *, $p < 0.05$ by Mann–Whitney U test. (F) The expression of genes shown in (E) does not significantly change over time in uninfected cells.

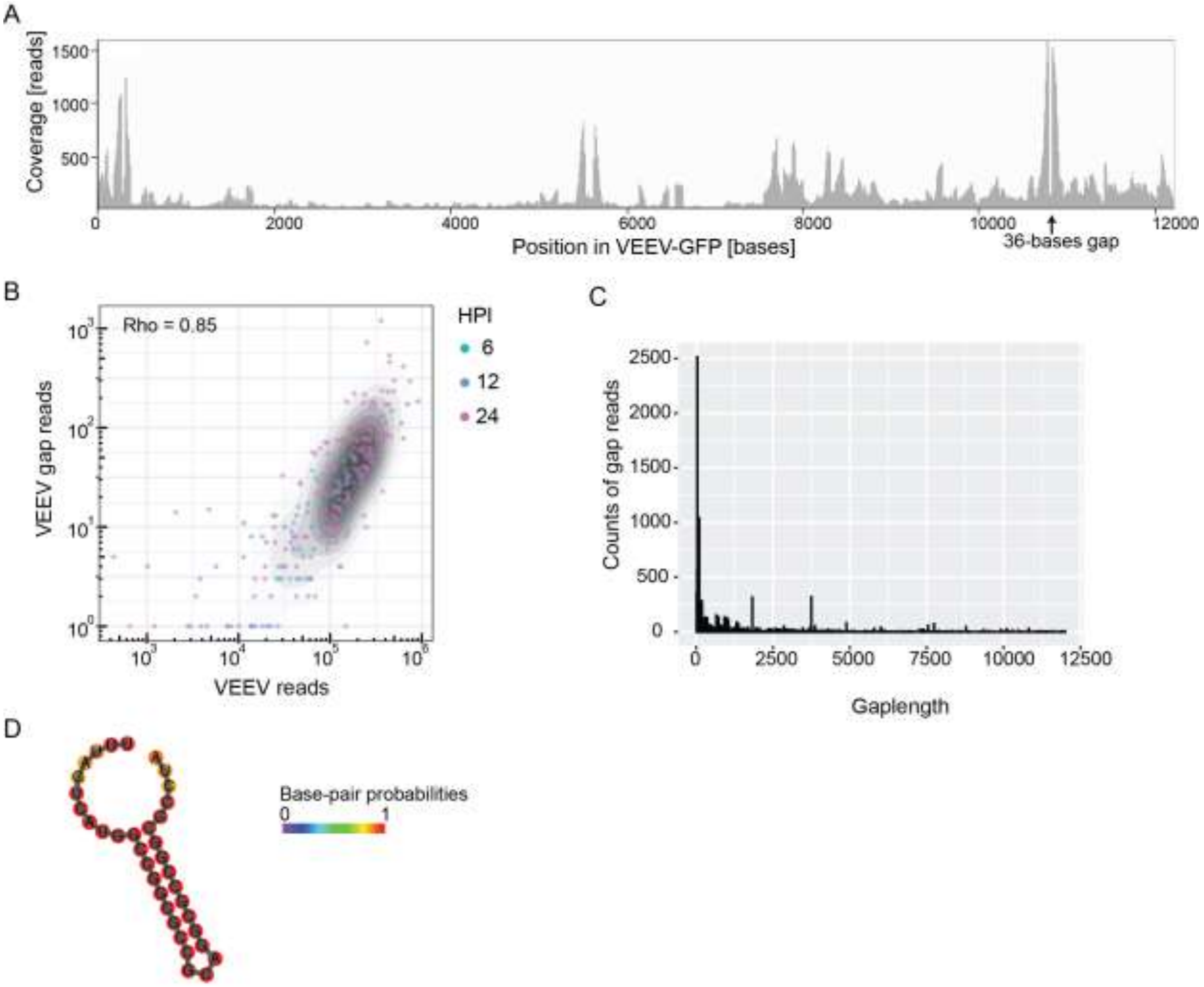


**S3 Fig. VEEV gap reads identified via viscRNA-Seq.** (A) Coverage of VEEV gap reads over the VEEV-TC-83-GFP genome. (B) Scatter plot of number of VEEV gap reads and VEEV total reads within cells with detected gap reads. Each dot represents a cell and colored by hpi. (C) Histogram of gap lengths indicating that the majority of gaps are shorter than 1000 nucleotides. (D) RNA structural prediction of the most common 36-base gap (arrow in A) via RNAfold web server. Scale bar indicates the possibilities of base pairing. Hpi, hours postinfection.

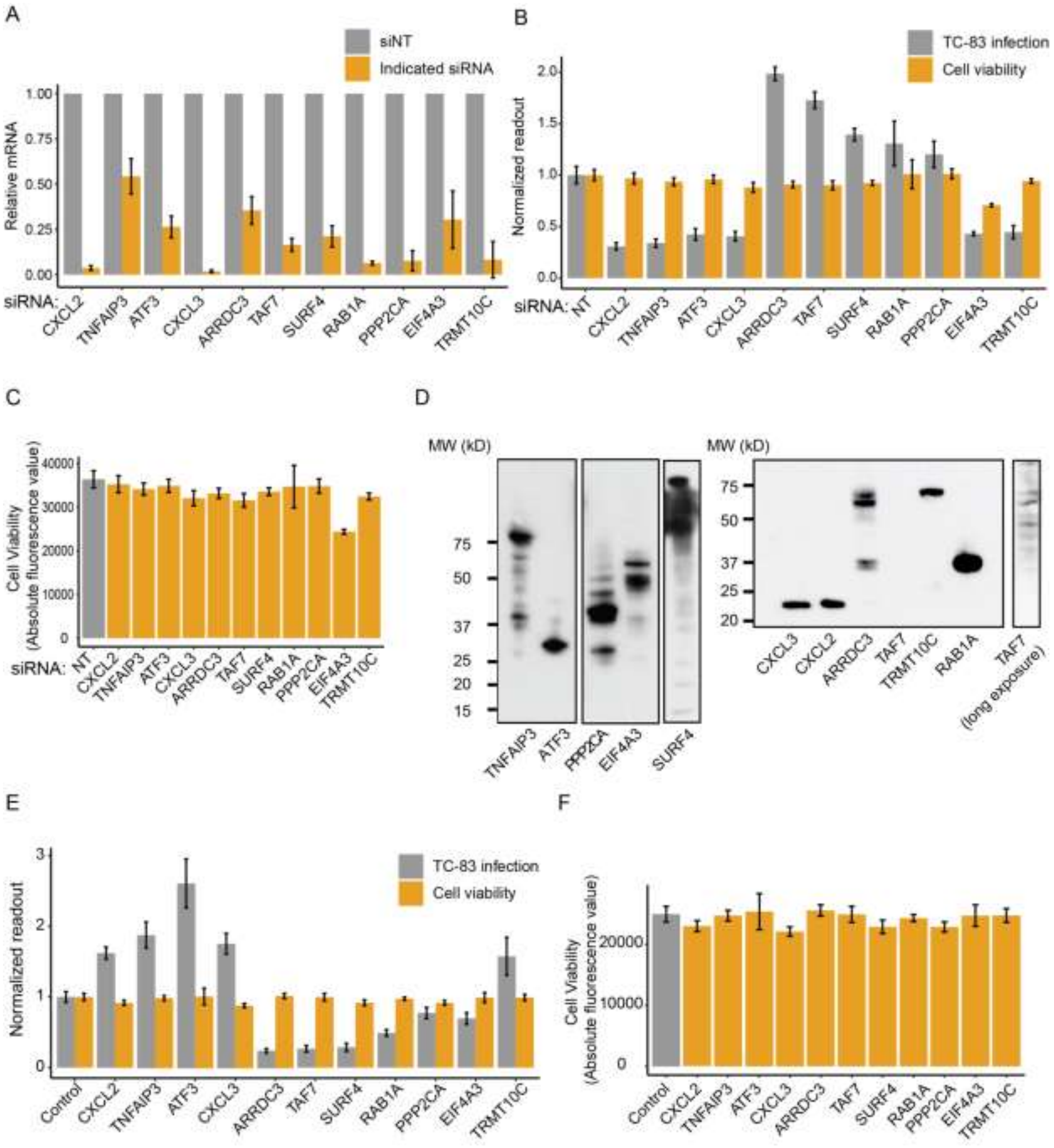


**S4 Fig. Loss-of-function and gain-of-function experiments for validation of candidate proviral and antiviral factors.** (A) Confirmation of gene expression knockdown in U-87 MG cells transfected with the indicated siRNAs or non-targeting control (NT) via qRT-PCR at 96 hours post-transfection. Results are relative to the level of the respective genes in the NT control. (B and E) Overall VEEV-TC-83 infection (grey) measured by luminescence assays and cell viability (orange) measured by alamarBlue assays in U-87 MG cells transfected with the indicated siRNAs (B) or ectopically expressing the indicated cellular factors (E) at 18 hpi with VEEV-TC-83-nLuc (MOI = 0.01). Data are expressed relative to siNT (B) or empty plasmid (E)

controls. (C and F) Absolute fluorescence values from the alamarBlue assays shown in B and E. (D) Confirmation of ectopic expression of the indicated gene products tagged with a FLAG-tag by Western blot in U-87 MG cells. Membranes were blotted with anti-FLAG antibody. Samples in the left panels were run on the same gel from which several lanes were cut out. TAF7 expression on the right is shown at a higher exposure. Data sets are pooled from two independent experiments with six replicates each (B,C, E and F). Shown are means ± SD.

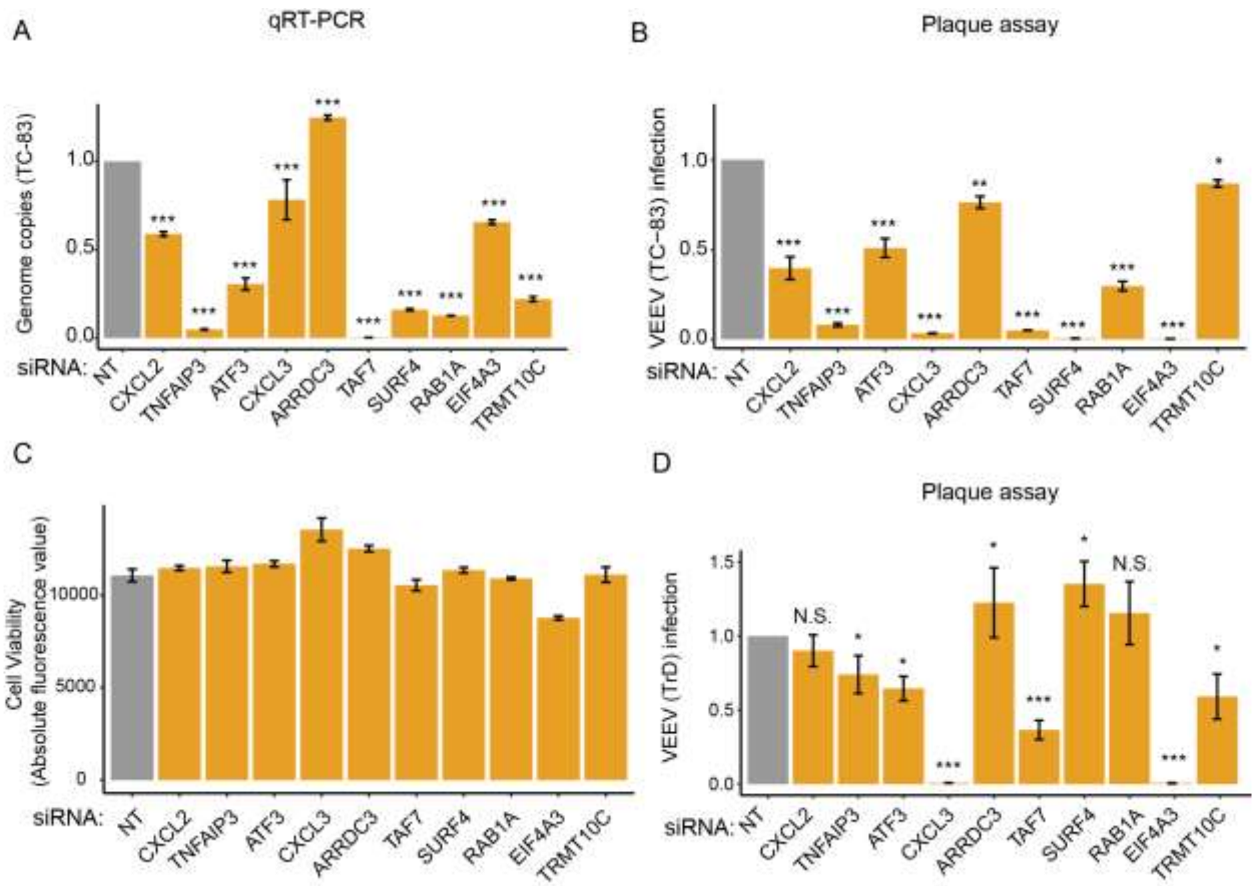


**S5 Fig. Functional relevance of viscRNA-Seq hits in cells infected with wild type TC-83 and TrD VEEV.** (A) Viral genome copies in lysates derived from U-87 MG cells transfected with the indicated siRNAs 24 hpi with non-reporter VEEV-TC-83 at an MOI of 0.01. (B and D) VEEV infection via plaque assays in U-87 MG cells transfected with the indicated siRNAs 24 hpi with non-reporter TC-83 (B) and TrD (D) (MOI = 0.001). C. Cell viability via alamarBlue assays in U-87 MG cells transfected with the indicated siRNAs. Shown are means ± SD. Data are plotted relative to non-targeting (NT) siRNA control (A, B and D). Representative experiments of at least two conducted are shown. *, $q < 0.05$; **, $q < 0.01$; *** $q < 0.001$ by 1-way ANOVA followed by False Discovery Rate (FDR) corrected multiple comparisons test. N.S, non-significant.

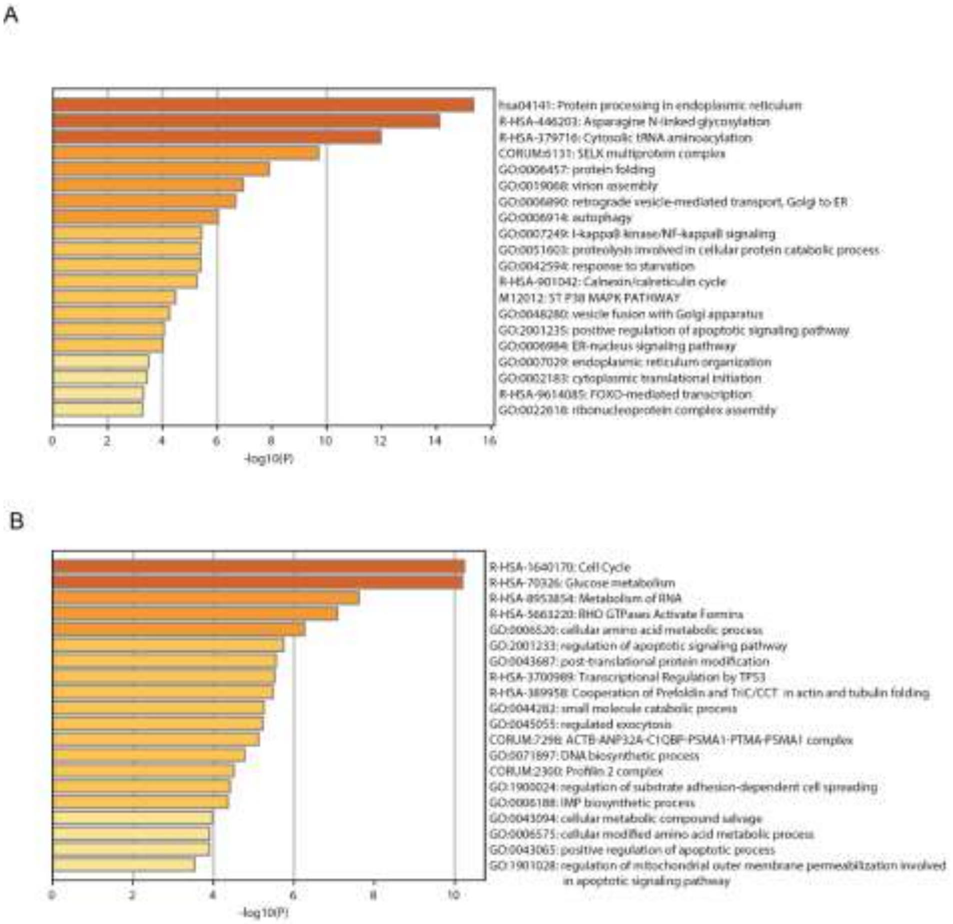


**S6 Fig. Pathway analysis for genes that positively correlated with VEEV 3'/5' read ratio and positively (A) or negatively (B) correlated with DENV and ZIKV.** Each bar represents a group of genes according to Gene Ontology, KEGG, or other databases of biological function. The plot was made using metascape.

**S1 Table.** VEEV capture oligonucleotides.

| Oligo name | Sequence (5’ to 3’) | Position on VEEV genome |
|---|---|---|
| VEEV_1 | AAGCAGTGGTATCAACGCAGAGTACTTCCTTATCAGTTATTTCCTTACAG | 353 - 377 |
| VEEV_2 | AAGCAGTGGTATCAACGCAGAGTACAGATAATTTTTCACTCTTGAGTACA | 1742 - 1766 |
| VEEV_3 | AAGCAGTGGTATCAACGCAGAGTACTTTTAGGTCTTATAATGGCTATGAG | 2442 - 2466 |
| VEEV_4 | AAGCAGTGGTATCAACGCAGAGTACTGCTGATAGTGATGGTATTTATATG | 3700 - 3724 |
| VEEV_5 | AAGCAGTGGTATCAACGCAGAGTACCTACTGACTTGTAATTGTTATCGTT | 4320 - 4344 |
| VEEV_6 | AAGCAGTGGTATCAACGCAGAGTACGTAGTAATTCTTCTTTTTCTTGGTC | 5823 - 5847 |
| VEEV_7 | AAGCAGTGGTATCAACGCAGAGTACTTCATTATTACACGCATATTTCTTG | 6383 - 6407 |
| VEEV_8 | AAGCAGTGGTATCAACGCAGAGTACGCATCTATAATCTTGACTTCCATAT | 7162 - 7286 |

**S2 Table.** siRNA sequences of candidate genes.

| Pool Catalog Number | Duplex Catalog Number | Gene Symbol | GENE ID | Gene Accession | GI Number | Sequence |
|---|---|---|---|---|---|---|
| L-007878-00 | J-007878-08 | CXCL3 | 2921 | NM_002090 | 54144649 | CCAACUGACAGGAGAGAAG |
| L-007878-00 | J-007878-09 | CXCL3 | 2921 | NM_002090 | 54144649 | UCCAAAGUGUGAAUGUAAG |
| L-007878-00 | J-007878-10 | CXCL3 | 2921 | NM_002090 | 54144649 | GGGAGACCAUAAUGUGUCA |
| L-007878-00 | J-007878-11 | CXCL3 | 2921 | NM_002090 | 54144649 | UACGAGGGUUCUACUUAUU |
| L-007877-02 | J-007877-07 | CXCL2 | 2920 | NM_002089 | 148298657 | CCGCAUCGCCCAUGGUUAA |
| L-007877-02 | J-007877-08 | CXCL2 | 2920 | NM_002089 | 148298657 | GAGCAGAGAGGUUUCGAUA |
| L-007877-02 | J-007877-09 | CXCL2 | 2920 | NM_002089 | 148298657 | GAUAGAAGGUUUGCAGAUA |
| L-007877-02 | J-007877-21 | CXCL2 | 2920 | NM_002089 | 148298657 | CAAUGUGACGGCAGGGAAA |
| L-009919-00 | J-009919-05 | TNFAIP3 | 7128 | NM_006290 | 26051241 | CUGCAGUACUUGCUUCAAA |
| L-009919-00 | J-009919-06 | TNFAIP3 | 7128 | NM_006290 | 26051241 | CAACUCAUCUCAUCAAUGC |
| L-009919-00 | J-009919-07 | TNFAIP3 | 7128 | NM_006290 | 26051241 | UCUGGUAGAUGAUUACUUU |
| L-009919-00 | J-009919-08 | TNFAIP3 | 7128 | NM_006290 | 26051241 | CAACGAAUGCUUUCAGUUC |
| L-008663-00 | J-008663-05 | ATF3 | 467 | NM_001030287 | 71902535 | GAGCUAAGCAGUCGUGGUA |
| L-008663-00 | J-008663-06 | ATF3 | 467 | NM_001030287 | 71902535 | GCAAAGUGCCGAAACAAGA |
| L-008663-00 | J-008663-07 | ATF3 | 467 | NM_001030287 | 71902535 | AGAAGCAGCAUUUGAUAUA |
| L-008663-00 | J-008663-08 | ATF3 | 467 | NM_001030287 | 71902535 | CGAGAAAGAAAUAAGAUUG |
| L-014063-01 | J-014063-09 | ARRDC3 | 57561 | NM_020801 | 32698735 | GAACGUUUGCUGCGAUAAA |
| L-014063-01 | J-014063-10 | ARRDC3 | 57561 | NM_020801 | 32698735 | UGCAAGAGGACAUGCGAAA |
| L-014063-01 | J-014063-11 | ARRDC3 | 57561 | NM_020801 | 32698735 | GUAUAAUCCGCGUGGAAUA |
| L-014063-01 | J-014063-12 | ARRDC3 | 57561 | NM_020801 | 32698735 | AGUCAGUGUAGCAUGAAUA |
| L-010622-01 | J-010622-09 | SURF4 | 6836 | NM_033161 | 19593984 | CCACAAGGGUAGUCGAACA |
| L-010622-01 | J-010622-10 | SURF4 | 6836 | NM_033161 | 19593984 | CGAAUAUUGGUAAGAUCGA |
| L-010622-01 | J-010622-11 | SURF4 | 6836 | NM_033161 | 19593984 | GCUCCCUGUUAGUGCCGUA |
| L-010622-01 | J-010622-12 | SURF4 | 6836 | NM_033161 | 19593984 | ACGUAUAUUUCAACGCCUU |
| L-008283-00 | J-008283-06 | RAB1A | 5861 | NM_004161 | 41350195 | CAGCAUGAAUCCCGAAUAU |
| L-008283-00 | J-008283-07 | RAB1A | 5861 | NM_004161 | 41350195 | GUAGAACAGUCUUUCAUGA |
| L-008283-00 | J-008283-08 | RAB1A | 5861 | NM_004161 | 41350195 | GGAAACCAGUGCUAAGAAU |
| L-008283-00 | J-008283-09 | RAB1A | 5861 | NM_004161 | 41350195 | UGAGAAGUCCAAUGUUAAA |
| L-003598-01 | J-003598-09 | PPP2CA | 5515 | NM_002715 | 57222566 | CCGGAAUGUAGUAACGAUU |
| L-003598-01 | J-003598-10 | PPP2CA | 5515 | NM_002715 | 57222566 | ACAUUAACACCUCGUGAAU |
| L-003598-01 | J-003598-11 | PPP2CA | 5515 | NM_002715 | 57222566 | UCAUGGAACUUGACGAUAC |
| L-003598-01 | J-003598-12 | PPP2CA | 5515 | NM_002715 | 57222566 | CAGGUAGAGCUUAAACUAA |
| L-020813-01 | J-020813-09 | TRMT10C | 54931 | NM_017819 | 8923404 | GAGAGUUAGUUAAACGGUA |
| L-020813-01 | J-020813-10 | TRMT10C | 54931 | NM_017819 | 8923404 | UUGACAUGGCUUACGAAAA |
| L-020813-01 | J-020813-11 | TRMT10C | 54931 | NM_017819 | 8923404 | GAGUUUAUCAACAGACUAA |
| L-020813-01 | J-020813-12 | TRMT10C | 54931 | NM_017819 | 8923404 | GGGAUAGGAAUAUGGACAU |
| L-020762-00 | J-020762-05 | EIF4A3 | 9775 | NM_014740 | 41327777 | UGACUAAAGUGGAAUUCGA |
| L-020762-00 | J-020762-06 | EIF4A3 | 9775 | NM_014740 | 41327777 | GGUGAAACGUGAUGAAUUG |
| L-020762-00 | J-020762-07 | EIF4A3 | 9775 | NM_014740 | 41327777 | GAUGAACGUUGCUGAUCUU |
| L-020762-00 | J-020762-08 | EIF4A3 | 9775 | NM_014740 | 41327777 | GAUAUGAUUCGUCGCAGAA |
| L-013669-00 | J-013669-05 | TAF7 | 6879 | NM_005642 | 14717406 | GAGAAGGGCAGUACAGUCU |
| L-013669-00 | J-013669-06 | TAF7 | 6879 | NM_005642 | 14717406 | GGACAAGCUAAAUGAAUCA |
| L-013669-00 | J-013669-07 | TAF7 | 6879 | NM_005642 | 14717406 | AAUCACUCCUAGAGAAGUA |
| L-013669-00 | J-013669-08 | TAF7 | 6879 | NM_005642 | 14717406 | UAGUAGACCUGCCCUGUGU |
| D-001810-10 | D-001810-01 | ON-TARGETplus Non-targeting Control | 0 | | 0 | UGGUUUACAUGUCGACUAA |
| D-001810-10 | D-001810-02 | ON-TARGETplus Non-targeting Control | 0 | | 0 | UGGUUUACAUGUUGUGUGA |
| D-001810-10 | D-001810-03 | ON-TARGETplus Non-targeting Control | 0 | | 0 | UGGUUUACAUGUUUUCUGA |
| D-001810-10 | D-001810-04 | ON-TARGETplus Non-targeting Control | 0 | | 0 | UGGUUUACAUGUUUUCCUA |

**S3 Table.** qRT-PCR primer sequences.

| NAME | SEQUENCE |
|---|---|
| CXCL2-F | GGCAGAAAGCTTGTCTCAACCC |
| CXCL2-R | CTCCTTCAGGAACAGCCACCAA |
| CXCL3-F | CGCCCAAACCGAAGTCATAG |
| CXCL3-R | GCTCCCCTTGTTCAGTATCTTTT |
| ATF3-F | CCTCTGCGCTGGAATCAGTC |
| ATF3-R | TTCTTTCTCGTCGCCTCTTTTT |
| TNFAIP3-F | TCCTCAGGCTTTGTATTTGAGC |
| TNFAIP3-R | TGTGTATCGGTGCATGGTTTTA |
| ARRDC3-F | ATGCAAGAGGACATGCGAAAG |
| ARRDC3-R | GTGGAAGCCTTCTTCGGAATTAT |
| TAF7-F | GAGTGGACCGTGTTCCATTGG |
| TAF7-R | CAACTGTGGATACAAGCATCTGA |
| SURF4-F | TGCCTACAGCATTTTATGGGAC |
| SURF4-R | TCTTCCCTTCAGAACGGGATT |
| RAB1A-F | AGATTAAAAAGCGAATGGGTCCC |
| RAB1A-R | GCTTGACTGGAGTGCTCTGAAT |
| PPP2CA-F | CAAAAGAATCCAACGTGCAAGAG |
| PPP2CA-R | CGTTCACGGTAACGAACCTT |
| EIF4A3-F | AAGGGAGAGATGTCATCGCAC |
| EIF4A3-R | GCTTGAGTTTCACGAACCTGA |
| TRMT10C-F | GGAAATGAAAGCAGCAGCAAGGGAAG |
| TRMT10C-R | AAACTGCATGGCCTGGGCAC |
| GAPDH-F | GGAGCGAGATCCCTCCAAAAT |
| GAPDH-R | GGCTGTTGTCATACTTCTCATGG |
| ACTB-F | CATGTACGTTGCTATCCAGGC |
| ACTB-R | CTCCTTAATGTCACGCACGAT |
